# Neural dynamics underlying the persistence of perseverative thought in depression and anxiety

**DOI:** 10.1101/2025.07.21.666019

**Authors:** Dale Zhou, Jeesung Ahn, David M. Lydon-Staley, Emily B. Falk, Dani S. Bassett, Ayelet Meron Ruscio

## Abstract

Depression and anxiety are characterized by transdiagnostic symptoms, including perseverative thought: a class of thoughts such as rumination and worry that are negative, repetitive, and difficult to control. These thoughts contribute to substantial distress, poor treatment response, and increased risk of relapse. What makes perseverative thoughts persevere? Using task-based fMRI, we compared how adults with no lifetime psychopathology and adults with major depressive disorder, generalized anxiety disorder, or both, engaged top-down control processes to switch from personally-relevant perseverative or neutral thought to a working memory task. For only adults with clinical depression or anxiety, stopping perseverative thought was associated with more probable frontoparietal deactivation and more frequent default-mode activation versus stopping neutral thought. Using network control theory, we identified key control points that lead to these activity dynamics. We found that clinical perseverative thought elicited less controlled activity in the anterior cingulate cortex relative to thoughts in adults with no lifetime psychopathology, and lower control energy correlated with greater depression severity. The occipital-temporal, lateral prefrontal, and insular cortices also used less control energy in clinical perseverative thought. Low energy is characteristic of attractor states in dynamical systems theory, deep channels wherein the flow of activity naturally settles, analogous to how a ball needs little energy to roll to the bottom of a bowl yet more energy to leave it. Entrenchment in attractors provides a computational perspective on why top-down control signals relate to the persistence of clinical perseverative thought. These insights advance our understanding of the dynamic processes of perseverative thought, paving the way for novel interventions for depression and anxiety.

## Introduction

Depression and anxiety are highly prevalent mental health conditions that frequently co-occur^1,2^, raising questions about transdiagnostic factors that may be implicated in their onset and maintenance. Two promising factors are worry and rumination^3,4^. Worry typically focuses on uncertain future threats^5^, whereas rumination typically involves dwelling on past or present concerns^6,7^. Although worry and rumination have historically been studied separately, they share a core process of perseverative thought: negative, repetitive, and dyscontrolled thinking that persists past an initial trigger, irrespective of the specific content or temporal focus of the thoughts^3,8,9^. A distinguishing feature of perseverative thought is its quality of being “sticky” or difficult to control, contributing to substantial distress, prolonged physiological arousal, and intrusions into daily tasks that impair functioning^10–15^. In generalized anxiety disorder (GAD) and major depressive disorder (MDD)—the two disorders most strongly associated with perseverative thought—these thoughts can become Sisyphean occupations, inescapable and repeating despite efforts to control them.

Several theoretical accounts have been proposed to explain problems of regulation in perseverative thought. One prominent account, the impaired disengagement hypothesis, proposes that repetitive negative thoughts arise from impaired neural signaling of the cognitive conflict needed to engage control processes that release attention from negative thoughts^16^. This hypothesis links perseverative thought to under-restrained brain regions in the default-mode network due to inactivity of regions in the frontoparietal and cingulo-opercular/salience networks that are typically active during cognitive control processes such as inhibiting unwanted thoughts, orienting attention to prioritized goals, and switching tasks^17–22^. Within the salience network, the anterior cingulate cortex (ACC) and anterior insula are engaged in tasks that require divided attention, executive control of automatic responses by signaling response conflict, and switching between large-scale network activation to deploy attention or working memory^23–30^. Consistent with the impaired disengagement hypothesis, ACC activity is robustly implicated in perseverative thought^31^, in which the dorsal ACC is associated with switching between default-mode and frontoparietal activation when cognitive control is needed, while the pregenual and subgenual ACC are associated with negative affect processing^32–34^. In contrast, the default-mode network is typically active during self-related thoughts, mind-wandering, autobiographical memory, and both future-oriented and past-oriented thinking^35^. Default-mode regions may be overly engaged in perseverative thought, rumination, and worry^31,36–45^, as well as in MDD and GAD^19,38,41,46–50^. Due to impaired top-down neural signaling, the default-mode network may be under-restrained and result in perseverative thought as a form of mind wandering focused on negative memories and self-concepts^31,42,46,51^.

Despite these conceptual and empirical advances, it remains puzzling why under-restrained mind wandering, in principle free to fluctuate and roam expansively, instead becomes “stuck” to highly repetitive thoughts^52–54^. A better understanding is needed of what dynamic processes underlie the difficulty of controlling the inherently dynamic process of perseverative thought. Several limitations of prior work can be addressed to make progress on this problem, including a tendency to infer maladaptive mind-wandering from abnormal resting-state neural activity without a corresponding experimental task^55,56^, a lack of a transdiagnostic approach to understanding shared and distinct processes of perseverative thought in worry versus rumination^57–60^, and a reliance on methods that do not study perseverative thought as a dynamic process^36,44,56,61,62^.

Dynamical brain measures can help adjudicate between competing explanations for uncontrollable perseverative thought. Two dynamic measures of brain activity, the transition probability and fractional occupancy, describe how spatiotemporal activity changes and recurs over time^63^. Transition probability refers to likelihood of moving from one neural activity pattern to another, whereas fractional occupancy captures the proportion of time spent in each activity pattern. Together, these measures offer insights into the cognitive inflexibility characteristic of persistent, repetitive thought^64^. Moreover, network control theory offers additional mathematical tools to understand how easy or difficult it is to change neural activity with minimal energy expenditure^65–69^. Treating brain function as a dynamical system, network control theory provides tools to model how activity evolves over time with a balance of spontaneous and controlled dynamics. Spontaneous dynamics spread activity across the network of anatomical connections with no external input, characterizing spontaneous thought that involves little effort^53,70^. Controlled dynamics are the result of various control points across the brain adding input from moment to moment, biasing how activity spreads according to target states and environmental feedback, characterizing more effortful processing^71–73^. Control energy is the integrated input across various control points within controlled dynamics and can be interpreted in terms of metabolic energy, mental effort, and transience^67,71,74,75^. Network control theory can be used to quantify the energy of top-down control from specific brain regions, like the ACC, as they bias activity in the transition away from perseverative thought toward a competing task^76^. There is evidence that higher-energy perturbations are more likely to foster more variable and transient rather than persistent cognitive states^73–75^. Persistence, by contrast, is a property of low-energy, stable dynamics^71^. This low energy is consistent with the proposed prominence of efficient, automatic processing when individuals face what are perceived as unattainable goals, according to the automatized information processing and energy minimization hypotheses of depression and anxiety^77,78^. These methods can advance our understanding of the entrenched nature of perseverative thought by shedding light on what types of thoughts and their brain processes may be more difficult to change or stop.

Here we use task-based fMRI to measure brain activity in 28 community-dwelling adults who were diagnosed with GAD or MDD (clinical group) and 16 adults with no lifetime mental disorder (non-clinical group) while they performed a novel thought-control task (**Figure 1**). During an initial thought period, personally-relevant perseverative or neutral thoughts were recalled using individualized autobiographical cues identified in an earlier semi-structured clinical interview. During a subsequent cognitive task period, participants were asked to stop the cued thoughts and perform a simple auditory 1-back task designed to be cognitively undemanding yet require sustained attention that competes with ongoing thought. Consistent with the impaired disengagement, automatized information processing, and energy minimization hypotheses, we hypothesize that compared to those without an anxiety or depression diagnosis, those with depression and anxiety show periods of worry and rumination that correspond to more frequent default-mode dynamics, less frequent frontoparietal dynamics, and lower-energy brain dynamics than in periods of neutral thought. Low-energy states are known as “attractors,” wherein the states are entrenched at a central location in an energy basin. The evolution of neural activity can be visualized as a ball rolling down into deepened dales along an energy landscape underlying activity patterns^79,80^. The trajectory of the ball models how the brain changes its activity to store and retrieve memory traces^81–84^ and switch between tasks^69,74,85^. Based on the known role of cognitive control regions in the inhibition of unwanted thoughts^25,86,87^, we hypothesize that the ACC and other regions in the frontoparietal and salience networks exhibit activity in a basin of attraction that enables reduced energy expenditure at the cost of becoming more entrenched in zones of self-referential and negative thought. The pull of such attractors may explain the persistence of perseverative thoughts in depression and anxiety^88–90^, reflecting a Sisyphean effort that would now be required to repeatedly push neural activity out of deepened energy basins. The combined neural dynamics approach can help us understand and potentially interrupt the cycle of repetitive negative thinking in depression and anxiety.

**Figure 1.**
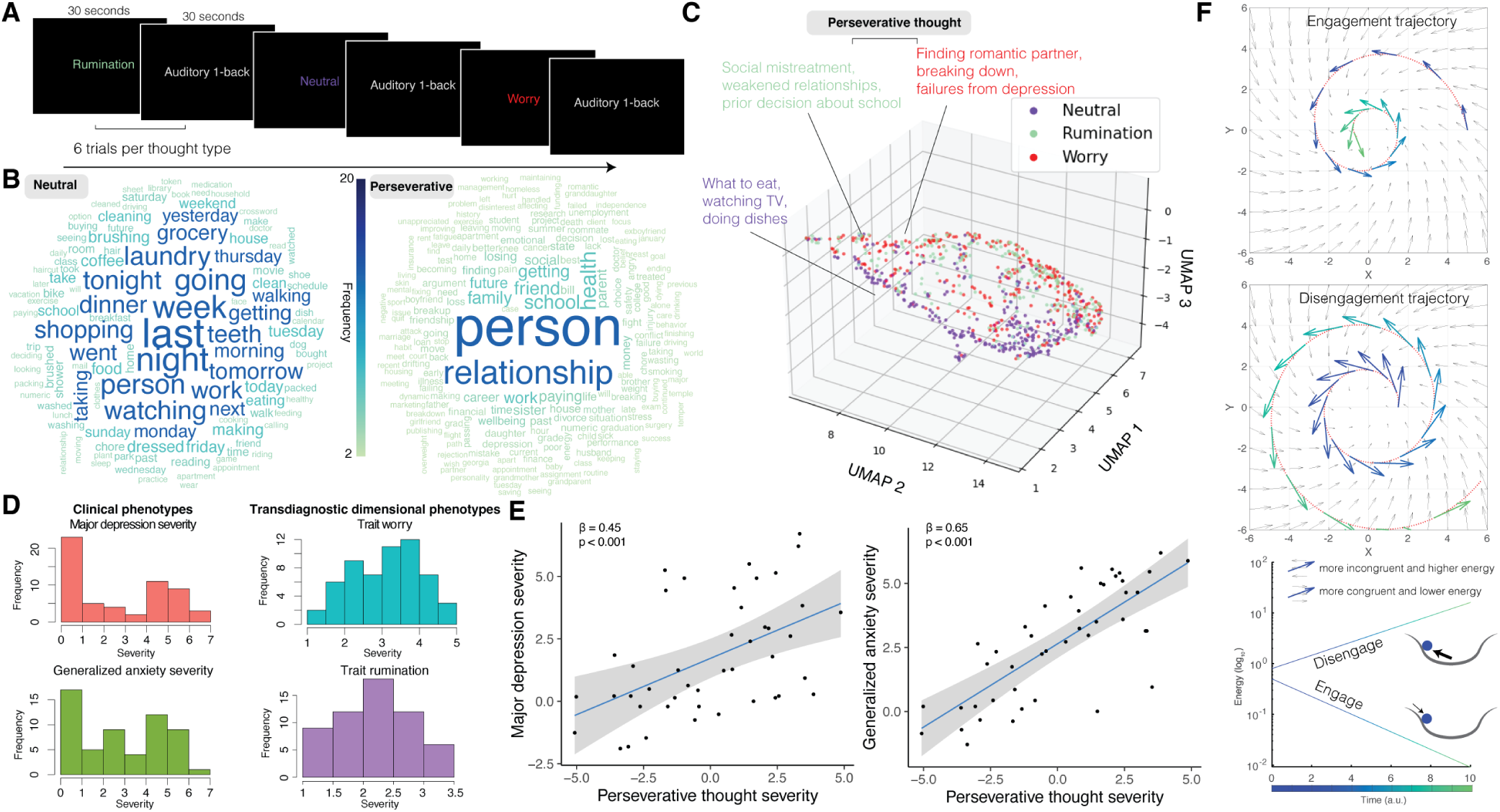
Task design, personalized stimuli, and perseverative thought. **(A)** The fMRI task instructed participants to transition from thinking about personally relevant worry, rumination, or neutral topics to performing an auditory 1-back working memory task. This design allowed us to evaluate the dynamic brain states involved in disengaging from perseverative thought (worry or rumination) and shifting to a different cognitive task. **(B)** Thought topic cues were individualized to participants based on a semi-structured interview and then presented in the scanner. To give a broad overview of the topics included, we visualize word frequency across the phrase stimuli. *Left*: neutral thought words. *Right*: perseverative thought words. Word size and color denote frequency across participants. Names have been replaced by “person.” **(C)** Each thought cue is a data point, and the distance between thoughts reflects the dissimilarity of the thought contents. Axes represent three primary dimensions of variation with arbitrary units determined by a Uniform Manifold Approximation and Projection embedding. Visually, neutral (purple) thoughts are distinguishable from rumination (green) and worry (red). Rumination and worry are each more heterogeneous in content than neutral thoughts. **(D)** Distribution of major depression severity, generalized anxiety severity, and transdiagnostic trait worry and rumination severity across all participants. Major depression and generalized anxiety severity represent clinical severity ratings assigned by interviewers on the Anxiety and Related Disorders Interview Schedule for the Diagnostic and Statistical Manual of Mental Disorders using a scale from 0 (none) to 8 (very severe). Worry severity represents participants’ mean score on the Penn State Worry Questionnaire, with a response scale from 1 (not at all typical) to 5 (very typical). Rumination severity represents participants’ mean score on the Ruminative Responses Scale, with a response scale of 1 (almost never) to 4 (almost always). **(E)** Major depression severity and generalized anxiety severity are strongly related to perseverative thought severity, underlining the usefulness of perseverative thought phenotypes for a dimensional approach to psychiatry. Data points indicate individuals and the grey ribbon indicates the 95% confidence interval from linear regression. **(F)** Hypothesized effect of attractors (vector field) on perseverative thought trajectories (red dotted line) over time (blue-to-green color). *Top*: Engaging perseverative thought involves moving inward congruently with the attractor field. *Middle*: Disengaging perseverative thought involves pushing outward against the attractor field. *Bottom*: Thought dynamics congruent with the attractor field use less energy than thought dynamics that must push against the attractor field, like a ball rolling into versus being pushed out of a basin.

## Results

### Challenges in disengaging default-mode states and engaging frontoparietal states after perseverative thought

Is brain state persistence greater during known periods of thought perseveration than during neutral thought blocks? To answer this question, six brain states were identified across all participants’ fMRI data using a *k*-means clustering approach previously shown to identify distinctive and interpretable recurring coactivation patterns across the brain **(Figure 2A-B; Supplementary Figure S1)**^63^. To aid interpretability, each brain state is referred to by the name of the large-scale brain network that is most similar to the positive and negative amplitudes of that brain state. Consequently, each state is associated with an activated and a deactivated brain network, including visual (VIS), somatomotor (SOM), frontoparietal (FPN), default-mode (DMN), ventral attention (VAT), and dorsal attention (DAT) areas^18^.

**Figure 2.**
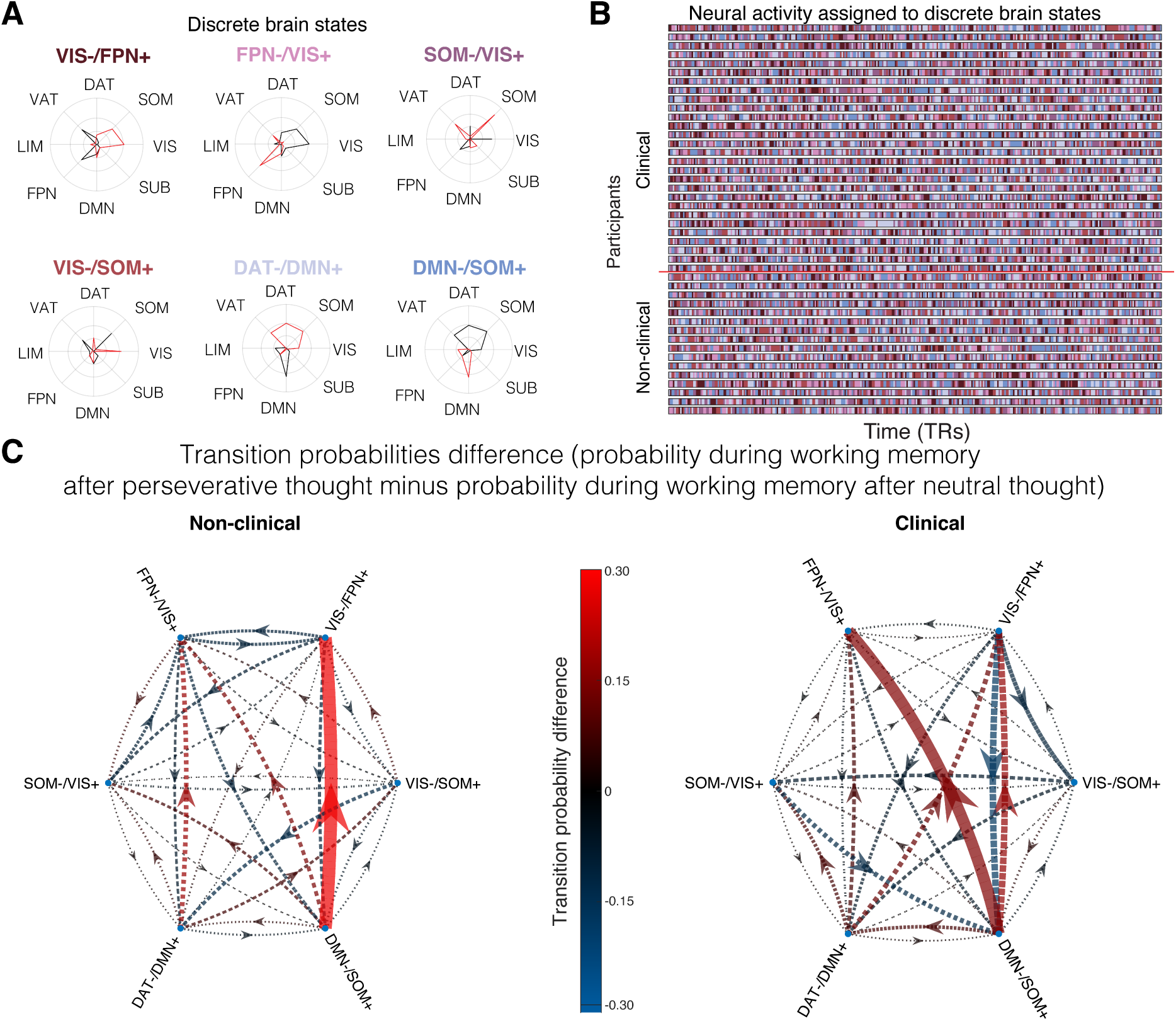
Default-mode and frontoparietal dynamics during the transition from perseverative thought to working memory. **(A)** Six centroids of fMRI activity were identified to characterize discrete brain states across the activity of all participants. For interpretability, each brain state is referred to by the name of the network of regions that is most prominent: visual (VIS), somatomotor (SOM), dorsal attention (DAT), ventral attention (VAT), limbic (LIM), frontoparietal (FPN), and default-mode (DMN). Radar plots depict which networks have more deactivation (red), denoted with a “-”, and which have more activation (black), denoted with a “+”. **(B)** The task activity of all participants was assigned to brain states. **(C)** Colors depict the difference in transition probability for the working memory task after perseverative thought versus after neutral thought. For the non-clinical group, neural activity during working memory had a higher probability of transitioning from deactivated default-mode network states (DMN-/SOM+) to more activated frontoparietal network states (VIS-/FPN+) after periods of perseverative thought than after neutral thought. For the clinical group, this higher probability of transitioning from default-mode network states (DMN-/SOM+) to frontoparietal network states (VIS-/FPN+) was only partially observed; instead there was a higher probability of transitioning from deactivated default-mode network states (DMN-/SOM+) to deactivated frontoparietal network states (FPN-/VIS+). This pattern is consistent with diminished engagement of cognitive control regions in the clinical group. Dashed arrows indicate transition probability differences that are not statistically significant, whereas solid arrows indicate significant differences between task conditions.

We first examined how the dynamic brain measures of transition probability (the likelihood of moving from one state to another; see **Methods**) and fractional occupancy (the proportion of time spent in each state) change during task performance. Given our particular interest in the ability to disengage from perseverative thought and transition to new cognitive states, we examined the transition probability during the working memory task when it followed worry or rumination compared to when it followed neutral thoughts (**Figure 2C**). Within the non-clinical group, the probability of transitioning from DMN-to FPN+ was greater when the working memory task followed perseverative thought (mean *M* = 30.31 ± 13.18%) versus neutral thought (*M* = 16.41 ± 13.05%; *t*(15) = 4.40, *p* = 0.02; all *p*-values adjusted for multiple comparisons using Bonferroni correction; see **Supplementary Figure S2** for FDR correction). These differences in transition probabilities are consistent with the need to disengage default-mode dynamics associated with self-relevant thoughts in order to engage frontoparietal dynamics associated with the goal-related processes of cognitive control. By contrast, within the clinical group we observed a greater probability of transitioning from DMN-to FPN-, instead of FPN+, when the working memory task followed perseverative thought (*M* = 30.98 ± 16.73%) versus neutral thought (*M* = 15.56 ± 16.43%; *t*(27) = 3.92, *p* = 0.02). These transition dynamics may indicate that, in the clinical group, disengagement from the DMN is preserved but engagement of FPN is impaired such that control processes are less responsive to the external demands of new tasks^16,91,92^.

FPN deactivation may be explained by its relationship with DMN activation, as these networks are thought to encompass dynamically opposing task-positive and task-negative regions, respectively. Indeed, within the clinical group but not the non-clinical group, the fractional occupancy of DAT-/DMN+ during the working memory task was greater when it followed perseverative thought (*M* = 0.21 ± 0.05) than when it followed neutral thought (*M* = 0.15 ± 0.06; *t*(53.58) = 3.84, *p* = 0.0003) (**Figure 3**; see **Supplementary Figure S3** for FDR correction). Across all participants, the fractional occupancy of DAT-/DMN+ during the working memory task following rumination was associated with greater trait rumination severity (*β* = 0.32, *p* = 0.03) and marginally with major depression severity (*β* = 0.77, *p* = 0.12). Increased occupancy of DAT-/DMN+ suggests that perseverative thought, more than neutral thought, evokes activity associated with less attention to shifting task demands and more mind-wandering and self-related thoughts.

**Figure 3.**
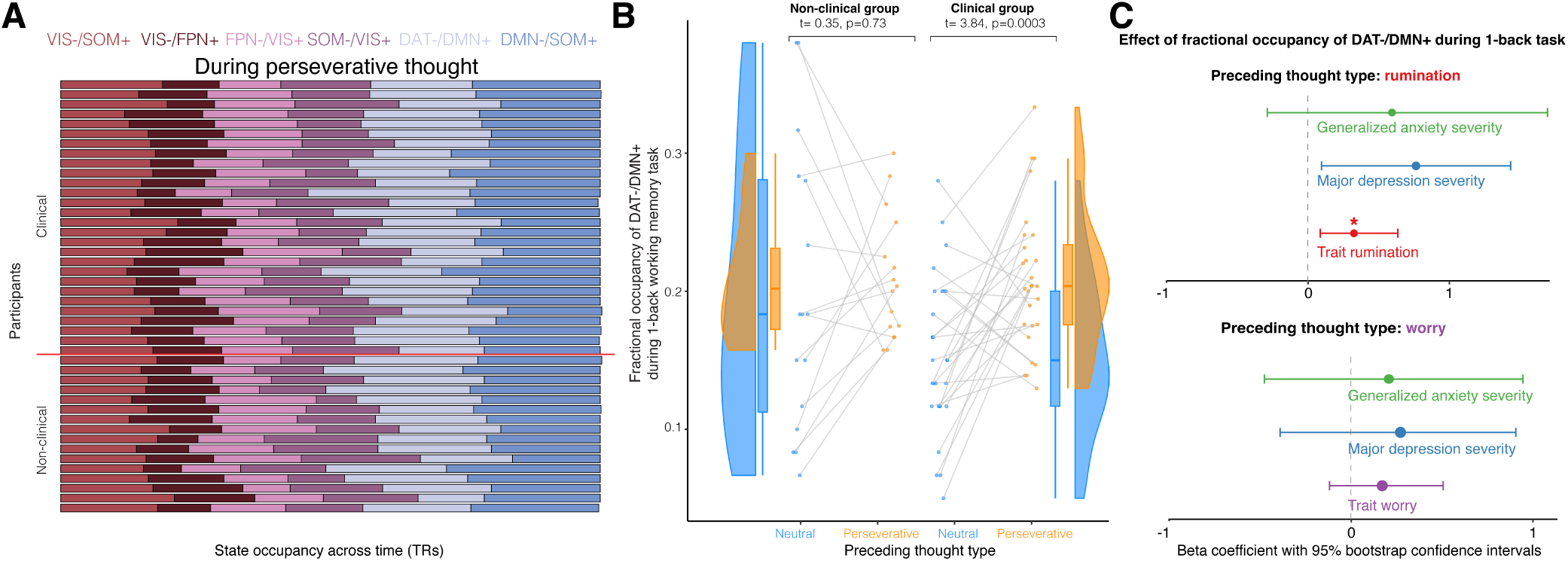
Default-mode fractional occupancies differ during working memory when transitioning from neutral thought versus perseverative thought. **(A)** Fractional occupancy of all brain states during perseverative thought (worry or rumination). **(B)** During the 1-back working memory task, participants in the clinical group occupied the DAT-/DMN+ state more when the task followed periods of perseverative thought than when the task followed neutral thought. This difference was not observed in the non-clinical group, nor for other brain states. (C) *Top:* When rumination preceded the working memory task, greater fractional occupancy of DAT-/DMN+ was associated with greater trait rumination severity and marginally with greater depression severity. *Bottom:* When worry preceded the working memory task, no effects were observed.

Despite these findings within each group, there were no significant differences in transition probability (all *t <* 2.97, all *p >* 0.18) or fractional occupancy (all *t <* 2.23, all *p >* 0.19) between groups, nor were there significant correlations between the other dynamic brain measures and dimensional measures of clinical symptoms (all *p*-values > 0.22; see **Supplementary Figure S4**). These negative findings motivate the application of network control theory tools that may be more sensitive to group differences and clinical measurements.

### Perseverative thought is associated with temporal sequences of brain activity entrenched in low-energy top-down signaling

We next applied measures of brain dynamics from network control theory to characterize the controlled disengagement from default-mode states to frontoparietal states following perseverative thought periods. We tested the role of the ACC in the dynamic control of brain activity during rumination and worry, analyzing them separately from perseverative thought to account for their distinct energy profiles. The control energy of a region is calculated as the energy contributed by that region to change all activity according to measured transitions between consecutive points of neural activity during the task paradigm **(Figure 4A-B)**. We then calculated the control energy used for thought types and their ensuing working memory task. We hypothesized that control energy is lower in the ACC for clinical compared to non-clinical perseverative thought, consistent with diminished cognitive control signals and a basin of attraction that compels repetitive and difficult-to-escape patterns of activity **(Figure 4C)**. Consistent with our hypothesis, perseverative thought periods were characterized by lower energy of ACC activity in the clinical group compared to the non-clinical group (**Figure 4D-E**). Across rumination trials, control energy was lower for participants in the clinical group (*M* = −0.30 ± 0.76) compared to the non-clinical group (*M* = 0.12 ± 0.84; *t*(171) = −3.95, *p* = 0.0007). A similar pattern was observed for worry, with lower control energy in the clinical group (*M* = −0.15 ± 0.87) than in the non-clinical group (*M* = 0.87 ± 0.92; *t*(171) = −3.24, *p* = 0.009). This group difference in control energy also held during the working memory task following rumination (clinical: *M* = −0.13 ± 0.87; non-clinical: *M* = 0.26 ± 1.04; *t*(171) = −3.10, *p* = 0.01); this pattern did not extend to worry (*t*(171) = −1.81*, p* = 0.43). These results suggest that the ACC used less energy when sending top-down signals to stop ruminating in the clinical group compared to the non-clinical group.

**Figure 4.**
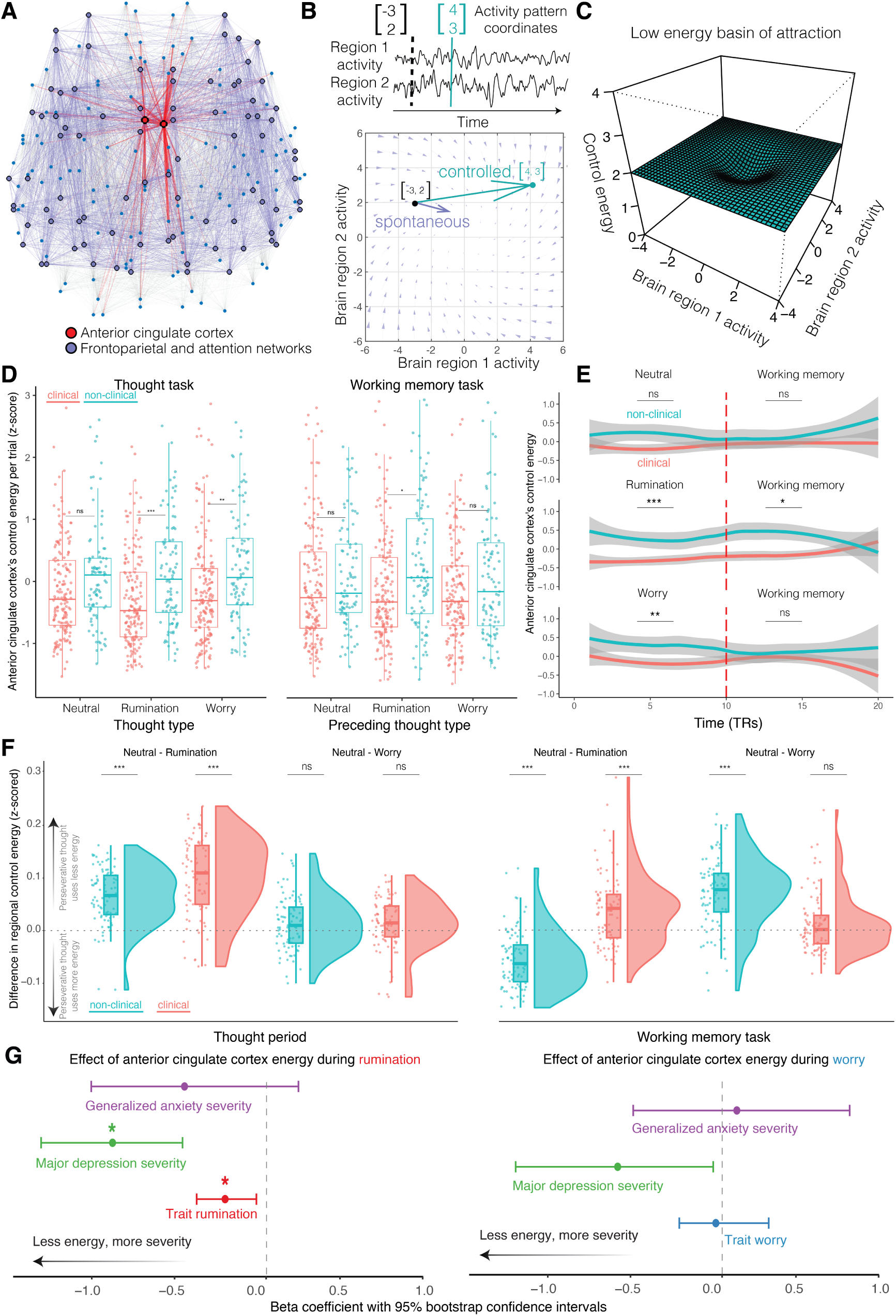
Ruminative thought occurs with brain states entrenched in low-energy top-down signaling. **(A)** The ACC (red) is embedded among larger frontoparietal and salience networks (blue) which send top-down signals during task performance via anatomical connectivity (edges) to the rest of the brain (nodes). **(B)** *Top*: The evolution of activity patterns can be modeled as movement in a dynamic neural state space. *Bottom*: Spontaneous activity (blue vector field) arises from a linear dynamics model of activity naturally flowing across structural connections over time. **Figure 4**. Spontaneous transitions are less energetically expensive (length of the blue arrow) than controlled activity (length of the cyan arrow), when control energy is required to counter the trajectory of spontaneous activity. **(C)** Low-energy transitions occur near basins of attraction, which are easier to linger in and more difficult to depart from. These low-energy dynamical properties may help explain the difficulty of disengaging from perseverative thought. **(D)** Trial-by-trial control energy during perseverative thought (rumination or worry) tends to be lower in the clinical group compared to the non-clinical group. Data points indicate trials split by clinical status and thought type. Control energy is averaged across regions in the frontoparietal, dorsal attention, and ventral attention networks. **(E)** As participants transition into the working memory task, only ruminative thought persists in low-energy activity in the ACC region of interest. **(F)** Each data point represents a region’s control energy during neutral thought versus either rumination (Neutral – Rumination) or worry (Neutral – Worry). Positive values indicate that rumination or worry used less energy than neutral thoughts, whereas negative values indicate that rumination or worry used more energy than neutral thoughts. Regional control energy within groups is lower for both clinical and non-clinical groups during rumination compared to the neutral thought period, but lower for only the clinical group during the subsequent working memory task. **(G)** *Left:* Participants who have lower ACC control energy during rumination periods tend to have greater trait rumination and depression symptom severity. *Right:* ACC control energy during worry periods is unrelated to symptom severity after controlling for multiple comparisons. ACC control energy has a trending relationship with depression symptom severity. Beta coefficients are from linear regressions controlling for age and sex. Brackets illustrate bootstrapped 95% confidence intervals. *: *p <* 0.05; **: *p <* 0.01, ***: *p <* 0.001 after Bonferroni correction.

Beyond the ACC, we also assessed network-level differences in control energy within groups. Given the known roles of cognitive control and attention in regulating spontaneous thoughts^70^, we perform paired *t*-tests comparing the average control energy across all 84 bilateral regions of the frontoparietal network, dorsal attention, and ventral attention networks between different pairs of thought types, using the Bonferroni method to correct for multiple comparisons (**Figure 4F**). We again found that rumination uses lower energy than neutral thought in both the clinical (*t*(83) = −12.60, *p <* 0.001) and non-clinical (*t*(83) = −11.33, *p <* 0.001) groups. We did not find differences in control energy between worry and neutral thought (both *t*(83) *<* −2.53, both *p >* 0.16). During the working memory task, we found that regions use less energy when transitioning from rumination than from neutral thought in the clinical group (*t*(83) = −5.32, *p <* 0.001), but use more energy when making this transition in the non-clinical group (*t*(83) = 9.03, *p <* 0.001). Regions use less control energy when transitioning from worry than from neutral thought in the non-clinical group (*t*(83) = 9.82, *p <* 0.001), but not in the clinical group (*t*(83) = −2.15*, p* = 0.41). These results suggest that top-down signals from the frontoparietal, dorsal attention, and ventral attention networks have lower-energy dynamics when stopping clinical rumination and higher-energy dynamics when stopping non-clinical rumination in transitioning to the working memory task.

Given these findings at the level of large-scale brain networks, we followed up our hypothesis-driven analysis of the ACC with a data-driven, exploratory analysis to examine differences in regional control energy within each brain region in the frontoparietal, dorsal attention, and ventral attention networks (the nodes that were modeled to transmit top-down control signals). Given a large number of comparisons for 145 lateralized regions, we use a less stringent FDR correction in this exploratory analysis. We found that the left occipital-temporal cortex had lower energy in the clinical group compared to the non-clinical group during perseverative thought (rumination: Wilcoxon statistic *W* (262) = 5859*, p* = 0.02; worry: *W* (262) = 6003*, p* = 0.02 **Supplementary Figures S5-S6**). Moreover, during rumination only, the left insula had lower energy in the clinical versus non-clinical group (*W* (262) = 5981*, p* = 0.02). Finally, during worry only, the right dorsal prefrontal cortex (*W* (262) = 6235*, p* = 0.04) and right lateral prefrontal cortex (*W* (262) = 6156*, p* = 0.03) had lower energy in the clinical group compared to the non-clinical group.

Lastly, we asked if lower energy is associated with the severity of perseverative thought. Given that biophysical systems tend towards energetic efficiency and repeated retrieval may strengthen and automatize memories in a Hebbian fashion, we reasoned that frequently repeated perseverative thought—like that in individuals with high trait rumination and worry, or with more severe symptoms of disorders associated with these traits—would be less costly to maintain and therefore more likely to persist. Consistent with this notion, we found that brain states with higher probability (Spearman’s *ρ*(262) = −.16, *p* = 0.009) and occupancy (Spearman’s *ρ*(262) = −.15, *p* = 0.017) tended to use less control energy during periods of perseverative thought, consistent with prior work finding that energetically efficient transitions are more frequent^69,75^. Lower control energy in the ACC during rumination was associated with greater trait rumination severity (*β* = −0.25, *p* = 0.028) and depression severity (*β* = −0.93, *p* = 0.026) but not generalized anxiety severity (*β* = −0.49, *p* = 0.44) **(Figure 4G)**. However, lower energy in the ACC during worry was unrelated to trait worry severity (*β* = 0.16, uncorrected *p* = 0.91), generalized anxiety severity (*β* = 0.14, uncorrected *p* = 0.68), or major depression severity (*β* = −0.58, uncorrected *p* = 0.12). Low-energy basins of attraction that support energy efficiency and persistence could explain why some perseverative thoughts persist and are difficult to disengage.

## Discussion

This study addressed how the dynamics of brain activity change when individuals are asked to stop repetitive negative thinking and transition to a different cognitive state. We used measures capturing the progression of brain states as individuals attempted to interrupt personally-relevant perseverative and neutral thought. Network control theory was used to identify key control points in the temporal progression of perseverative thought. We identified the ACC as well as the insula, dorsal prefrontal cortex, lateral prefrontal cortex, and occipital-temporal cortex as key control points for transmitting top-down signals to transition away from perseverative thought. Compared to neutral thought, perseverative thought corresponded to brain activity with greater frontoparietal deactivation, greater default-mode occupancy, and lower control energy. Control energy for both rumination and worry was lower in the clinical group than the non-clinical group; these group differences were not observed for neutral thought. Furthermore, this lower energy persisted beyond the rumination thought period, despite instructions to stop the thoughts and shift to a competing cognitive task. Our results suggest that perseverative thought, and rumination in particular, corresponds to brain activity that dwells in deepened dales, shedding new light on why those states are difficult to terminate, especially for depressed and anxious individuals.

Our findings contribute to a growing understanding of why thoughts perseverate by investigating their dynamic, temporal nature^79,93–100^. Examining perseverative thought as a dynamic process has advantages over static approaches in understanding how thoughts unfold over time^100–102^. Our findings support the idea that the persistent nature of perseverative thought—at the neural, as well as behavioral, level—corresponds to greater clinical severity. We found that the severity of trait rumination and major depression related to control signals with lower energy. Moreover, the fractional occupancy of default-mode network activation and dorsal attention network deactivation related to trait rumination severity. Some of our findings appear specific to rumination and depression rather than shared with worry and GAD. That distinction may relate to rumination’s focus on past experiences, engaging more control of the memory processes of large association regions that receive top-down control inputs.

Understanding the potential roles of cognitive control processes, including the detection of cognitive conflict, task-switching, and behavioral inhibition, is key to elucidating how to let go of perseverative thought^103–108^. Prior work found that highly connected hub regions, like the ACC, have strong recurrent connectivity related to low control energy^109^, characterizing how feedback circuits generate attractor states that support persistent, stable memory traces^84^. Here, we extend that work to perseverative thought, showing that neural activity during rumination and worry evolves in a deepened attractor basin in dynamical systems theory. The width and depth of attractors may be increased by overgeneralized experiences and repeated occupancy^96,110,111^. Over time, memory decontexualization from repeated retrieval may make rumination easier to engage, “stickier”, and more difficult to stop^112–114^. Our network control theory results are also consistent with the role of the ACC proposed by the impaired disengagement hypothesis. The ACC is involved in emotion processing and cognitive control^16^. Specifically, the ACC signals cognitive conflict to dorsal and lateral prefrontal regions, which facilitate the control of attention away from perseverative thought by biasing the activity of other regions^25^, including occipital-temporal regions associated with inhibiting old memories^115,116^ involving social and emotional content^117,118^. Our results add that a putative failure of control could also be viewed functionally as an energetically efficient allocation of control resources to salient yet unresolvable goal discrepancies^29,41,119,120^, with the consequence that perseverative thought gravitate toward and become stuck near dynamical attractors, producing inflexible and rigid activity^36,121^. Worry and rumination are ubiquitous experiences and can help people to anticipate threats, pursue or let go of frustrated goals, and process troubling, ambiguous, or traumatic events^78,111,122–125^. For these purposes, persistent engagement benefits from greater energy efficiency. Attractor states in neural activity provide promising physical and dynamical operationalizations to concepts from psychological control theories that regard negative thoughts, including perseverative thoughts, as efficient and automatized responses to salient and unresolved goal discrepancies^77,78,124,126^.

### Clinical implications

Our findings have implications for the treatment of perseverative thought and associated disorders. The library of network control theory tools could provide a dynamical systems approach that helps tailor interventions to specific patterns of vulnerability^68,96,127^. Understanding the temporal dynamics of perseverative thought at both neural and behavioral levels provides a foundation for designing interventions that aim to improve cognitive flexibility and emotion regulation. For instance, there is accumulating evidence that intervening on key control points, whether by using cognitive strategies^73^, neural stimulation^128^, pharmacological intervention^129^, or neurofeedback^130,131^, is associated with improved self-regulation, learning, and threat processing.

Network control theory may be especially useful for testing hypothesized effects of pharmacological agents on brain receptors in depressive and anxiety disorders^69,76^. Serotonergic drugs have been found to reconfigure neural connectivity^132^ and to flatten the energy landscape underlying neural activity^133,134^. Flattening the energy landscape makes attractor basins more shallow and weakens their pull on the system’s dynamics, providing a possible mechanism for why selective serotonin reuptake inhibitors have relevance in treating major depression and anxiety. In addition, benzodiazepines like alprazolam (Xanax) have anxiolytic effects through enhanced GABAergic inhibition^135^. Alprazolam was found to increase the energy cost of persisting in a neural state during recall of threatening stimuli and emotion identification, consistent with promoting more activity change and transient states by making persistence more costly^129^. The increase in control energy with alprazolam administration is consistent with the notion that energy helps shift brain activity for engaging in new tasks^69,74,85^ and that GABAergic perturbations can reduce the over-stability of attractors^136^.

The role of attractors in perseverative thought is also consistent with the success of psychotherapies focused on repeated practice of new cognitive and behavioral patterns as an alternative to automatic, habitual perseverative thought^15,137,138^. Such therapies include shifting from abstract to concrete and detailed modes of processing, from internal to external focus, and from passive thinking to active problem solving, to help disengage from entrenched associative memories and narratives^139–141^. The effort to control thoughts and delve into the past to problem-solve can further entrench their perseverance if the goals and plans to attain them are unworkable^140^. Mindfulness interventions have reduced rumination, worry, major depression, and generalized anxiety severity^142–145^. Their efficacy is supported in part by explicitly training ongoing thoughts away from abstract levels of processing and into a more concrete mode of processing, as well as by training a non-judgmental acknowledgment of transient thoughts and feelings rather than attempting to appraise, solve, or inhibit them^146,147^. Prior work applying network control theory found that mindful attention was associated with improved self-regulation, achieved by increasing the energy of top-down signaling to facilitate transitions to new brain states and counteract the pull of attractors^73^. Interestingly, greater control energy was related to faster intrinsic timescales of neural activity in association regions implicated in the abstract processing mode, more similar to the faster rates typically observed in sensorimotor areas associated with a concrete processing mode^148^. Faster intrinsic timescales of neural activity characterizes dynamics whereby past activity updates to present activity more rapidly, providing a neurophysiological signature for “being present.” Faster intrinsic timescales also characterize dynamics whereby the present neural activity is more distinct from recent activity, providing a neurophysiological signature for creating psychological distance from transient thoughts and feelings. By contrast, dynamical signatures of stability and slowness in mood changes were found to precede depressive states^94^, reflecting activity dynamics more characteristic of association areas implicated in abstract thought.

Finally, the role of attractors in perseverative thought is consistent with the success of therapies that promote disengagement from more automatic self-focused thought to external focus. For example, social connections helps individuals to naturally synchronize with and predict another’s thoughts, feelings, and actions, processes that blur the boundary between self and other ^149, 150^. These social ties, in addition to forms of external focus, such as engaging in enjoyable activities and spending time in nature^151,152^, may help propel the brain to new neural states that are more distant from attractors^42,153,154^. This collection of results begins to elucidate a dynamical systems framework whereby pharmacological and psychosocial therapies can help the brain disengage from entrenched attractor dynamics to improve clinical outcomes.

### Limitations and future directions

The present results should be considered in the context of several strengths and limitations. Strengths include a clinical sample assessed using gold-standard diagnostic and severity measures, personally relevant perseverative and neutral thought stimuli, and an ecologically valid experimental paradigm designed to capture what happens when individuals attempt to stop perseverative thought and shift to other activities requiring their attention. One limitation was that the non-clinical group had a modest size of 16 participants, so within-group statistical results should be interpreted with caution. Another limitation was that network control theory simulations were performed on an average structural network template because diffusion imaging data were not available in the present study. This strategy has been used successfully in prior psychiatric research when structural connectivity data were absent^129^. The control energy results in the present paper therefore focus on the effects of different activity targets on control energy, given an average of shared structural connectivity. However, depressed individuals differ in structural connectivity strength from nondepressed individuals in specific large-scale networks^155^. Neglecting to model differences in structural connectivity may lead to underestimation of differences between the clinical and non-clinical group and underestimation of differences between thought periods within groups that engage the differently affected large-scale network connections. Future studies should examine the effect of individual differences in structural connectivity on control energy or use different system identification methods to estimate transition dynamics in place of anatomical connectivity^156,157^. Additionally, our analysis assumes linear dynamics, a useful approximation that could be further refined to identify the structure of attractors using non-linear methods^99,156,158,159^.

We asked participants to identify distinct thought cues for rumination, worry, and neutral categories. Although this is a standard approach for isolating characteristics of each thought type, it fails to consider interactions between thought types, such as the possibility that rumination may progress to worry over time^160^. Furthermore, we explored low control energy as an explanation for the uncontrollability of thoughts, but other properties of thoughts—such as their valence and vividness—are also theorized to contribute to adverse outcomes^111,137^. Future work should examine how control energy relates to the positive or negative valence of thoughts, drawing on recent applications of network control theory to emotional processing in structural connectivity networks^161,162^ and in interrelated affective symptom networks^127,163–166^. Future work could also investigate how attractors widen and deepen to affect neural^74,159,167^, thought^51,168,169^, and emotion dynamics^53,170^. One possibility is that the width of attractors, or the distance of their pull, is related to abstract goal construal or over-general memory that lead the dynamics into the same space via multiple pathways. The depth of attractors, or their “stickiness,” could be related to the strength of meta-cognitive beliefs about the helpfulness of rumination or worry^171,172^, a paucity of alternative coping strategies, or a reliance on reactive versus proactive control that attempts to control a thought after already retrieving and further strengthening the memory^173^. Each may lead to more frequent and automatized deployment of perseverative thought that digs deeper dales. Lastly, future work could investigate how control energy relates to subjective feelings of controllability, either in the moment or when assessed in daily life.

## Conclusion

This work advances understanding of the temporal dynamics of perseverative thought by using network control theory to elucidate how the persistence of repetitive negative thoughts relates to neural activity in low-energy basins of attraction. The pull of such attractors relates to the persistence of perseverative thoughts in depression and anxiety. “Being stuck in a hole” is a metaphor that is core to therapeutic interventions to elucidate the pull of perseverative thought in depression and anxiety^140^. Our findings make this clinical metaphor more concrete: in perseverative thought, a Sisyphean effort would be needed to repeatedly push neural activity out of deepened dales. Our results illustrate that moving beyond static models of cognition, and examining why perseverative thought persists, is a critical step toward disrupting the cycle of negative thinking that is implicated in so many mental health conditions.

## Methods

### Participants

Adults ages 18 years or older were recruited from the Philadelphia community through electronic and print advertisements. Those who met inclusion criteria based on online symptom questionnaires were contacted and screened for fMRI exclusion criteria. Individuals who passed this two-phase screening process were invited to the laboratory, where final eligibility was determined using the Anxiety Disorder Interview Schedule for DSM-IV—Lifetime version (ADIS)^174^. Participants receiving a current, principal (most severe) diagnosis of GAD or MDD were eligible for the clinical group. Participants with no current and lifetime psychopathology, and no more than moderate levels of trait worry, were eligible for the non-clinical group. Interviews were administered by Master’s– and Bachelor’s-level diagnosticians who were trained to a high level of interrater agreement with the supervising licensed clinical psychologist. Diagnoses and clinical severity ratings were finalized by the full assessment team following discussion of each case. MDD and GAD severity were rated by interviewers on the ADIS using a scale from 0 (none) to 8 (very severe). Interrater reliability was high for GAD (*K* = 1.00) and MDD (*K* = 0.88) diagnoses based on blind, independent ratings of recorded interviews (*n* = 32) that were selected at random from ongoing studies with these populations. Worry severity represents participants’ mean score on the Penn State Worry Questionnaire, with a response scale from 1 (not at all typical) to 5 (very typical)^175^. Rumination severity represents participants’ mean score on the Ruminative Responses Scale, with a response scale of 1 (almost never) to 4 (almost always)^176^. Perseverative thought severity is the mean of the *z*-scaled worry and rumination severity ratings.

All participants were right-handed, spoke and read English fluently, and had normal or corrected-to-normal vision. Standard MRI exclusion criteria were applied, including metal in the body, claustrophobia, and pregnancy. To reduce potential confounds, we also excluded individuals with neurological disorders, a history of head injury, or current use of psychoactive medications other than antidepressants. Antidepressant medications were permitted in the clinical group to enhance external validity, but only on a stable dosage. To further enhance external validity, we permitted comorbid mental disorders as long as GAD or MDD was the principal disorder. However, individuals with active suicidal intent, acute psychosis, or a current substance use disorder were excluded from participating and connected with clinical referrals.

Of the initial sample of 54 participants who completed the study, seven (3 GAD, 3 MDD, 1 non-clinical) were excluded from analyses due to excessive movement during scanning. The final sample included 30 clinical participants, of whom 13 (43%) had GAD only, 15 (50%) had MDD only, and 2 (7%) had both conditions. Finally, 2 participants in the clinical group were excluded for poor quality and incomplete data.

The analyzed sample included 28 participants in the clinical group (31.21 ± 10.18 years old; 18 female, 10 male) and 16 participants in the non-clinical group (27.94 ± 9.64 years old; 11 female, 5 male). The clinical group did not differ significantly from the non-clinical (*n* = 17) group in sex (*χ*^2^(3) = 5.80, *p* = 0.12), age (*t*(32.81) = –1.06, *p* = 0.30), or years of education (*χ*^2^(12) = 14.9, *p* = 0.25).

### Worry and rumination interview for in-scanner thought cues

A semi-structured interview was used to elicit the specific thought topics that were currently most relevant for each participant and would continue to be relevant during the coming week. Separate sections probed worry (“worries that you’ve been having about bad things that might happen in the future”) and rumination (“thinking or talking to yourself about bad things that have happened, including dwelling on how you feel and what has made you feel this way”). A parallel set of questions probed neutral thoughts (“thoughts about day-to-day things that are neither positive nor negative, and that do not stir up strong feelings for you”). These personally relevant, verbal-linguistic neutral thoughts, elicited using assessment procedures identical to those for worry and rumination, served as conservative comparison stimuli for perseverative thought. Participants rated the typical intensity, uncontrollability, and negative and positive affect associated with each thought topic on separate 0 (not at all) to 8 (extremely) Likert-type scales. Based on these ratings, participants selected their six most salient worry, rumination, and neutral thought topics; three neutral topics were future-oriented and three were past-oriented to match the temporal orientation of worry and rumination, respectively. Participants then generated a short phrase (3-4 words) that would be used to cue each thought topic in the scanner.

### Perseverative thought task paradigm

Within one week of their interview, participants were scanned while completing an experimental paradigm, developed for this study, to probe the neural substrates of intentional and uncontrolled perseverative thought. Each trial began with a cued thought block. A cue phrase evoking a worried, ruminative, or neutral thought topic—generated in the earlier interview—was displayed on a screen for the duration of the block. Participants were instructed to think about the cued topic as intensely as they could, in the way they normally thought about it, until they were asked to stop. Similar instructions have been used successfully in prior behavioral studies to induce worry and rumination^160,177^. A block length of 30 seconds was used based on pilot testing showing that this interval reliably evokes an intense, sustained period of induced thought, even in non-clinical samples.

After 30 seconds, the cue phrase disappeared from the screen, and participants were instructed to stop thinking about the cued topic and focus all of their attention on a cognitive task. In this n-back task, auditory stimuli (nonsense sounds) were presented one at a time over headphones. Participants wore the headphones for the entirety of the experiment. Participants indicated via key press whether each sound was the same as the sound that immediately preceded it. Stimuli were presented every 3 seconds over a block of 30 seconds.

The n-back task was chosen because it engages top-down brain networks associated differentiable from the more automatic processing involved in worry or rumination^178,179^. The task does not have substantial practice effects, as accuracy was already near ceiling. The 1-back task requires cognitive engagement but is minimally demanding, allowing a greater possibility of task-unrelated thoughts to intrude^180^ and is repeatable throughout the experiment. Moreover, the task stimuli are emotionally neutral, so would not be expected to overlap with or evoke perseverative thought. Taken together, the task offered an ecologically valid measure of the ability to control unwanted thoughts.

Out of the scanner, participants were debriefed about their experiences during the study. They then engaged in relaxed breathing with pleasant imagery to dispel any negative emotion, and were compensated for their time.

### MRI data acquisition

Neuroimaging was conducted using a 3-T Siemens Tim Trio MRI (Siemens Medical Systems, Iselin, NJ) with T1-weighted MPRAGE acquisition (repetition time = 1620 msec; echo time = 3.09 msec; field of view = 192 × 256 × 160 mm; voxel dimensions = 1 × 1 × 1 mm; 160 slices) and gradient echo T2*-weighted echoplanar images, acquired using an optimized pulse sequence (repetition time = 3000 msec; echo time = 30 msec; field of view = 64 × 75 × 63 mm; voxel dimensions= 2 × 2 × 2 mm; 49 interleaved slices). Task data were collected in 3 runs, each containing 6 blocks (160 images * TR (3 sec) = 8 min). Participants were assigned to either condition A or B with counterbalanced block orders.

### fMRI preprocessing

To ensure the reproducibility of data processing and analysis, brain image data (task-based functional MRI scans, T1– and T2-weighted structural MRI scans; *n* = 47) were organized in the standardized Brain Imaging Data Structure (BIDS;^181^) format using HeuDiConv (Version 0.8.0;^182^). The task-based fMRI scans were then preprocessed using fMRIPrep (Version 20.0.6;^183^), which is based on Nipype (Version 1.4.2;^184^). A reference volume and its skull-stripped version were generated using a custom methodology of fMRIPrep. A B0-nonuniformity map (or fieldmap) was estimated based on two echo-planar imaging (EPI) references with opposing phase-encoding directions, with 3dQwarp^185^ with AFNI 20160207.

Based on the estimated susceptibility distortion, a corrected EPI reference was calculated for a more accurate co-registration with the anatomical reference. The BOLD reference was then co-registered to the T1w reference using bbregister from FreeSurfer, which implements boundary-based registration^186^. Co-registration was configured with six degrees of freedom. Head-motion parameters with respect to the BOLD reference (transformation matrices, and six corresponding rotation and translation parameters) were estimated before any spatiotemporal filtering using mcflirt (FSL Version 5.0.9^187^). BOLD runs were slice-time corrected using 3dTshift from AFNI 20160207^185^.

The BOLD time-series were resampled onto their original, native space by applying a single, composite transform to correct for head motion and susceptibility distortions. The BOLD time-series were resampled into standard space, generating a preprocessed BOLD run in MNI152NLin2009cAsym space. All resamplings were performed with a single interpolation step by composing all of the pertinent transformations (i.e., head-motion transform matrices, susceptibility distortion correction when available, and coregistrations to anatomical and output spaces). Gridded (volumetric) resamplings were performed using antsApplyTransforms (ANTs), configured with Lanczos interpolation to minimize the smoothing effects of other kernels^188^. Non-gridded (surface) resamplings were performed using mri_vol2surf (FreeSurfer). Various confounds (e.g., framewise displacement, DVARS, global signal) were also calculated for each TR and logged in a confounds file. The outputs from fMRIPrep were then manually checked for quality to ensure adequate preprocessing.

### fMRI time series extraction

We extracted task-based BOLD time-series data for each participant using Nilearn’s NiftiLabelsMasker (version 0.7.1^189^) in the following order. To begin, BOLD signals were detrended and temporal filters were applied to retain signals in the range of 0.01–0.10 Hz. Signals were denoised by removing 36 confounds orthogonally to the temporal filters. Confounds were generated from the fMRIPrep preprocessing procedure, which included six realignment parameters, mean signal in white matter, CSF and mean global signal, as well as the first power and quadratic expansions of their temporal derivatives. Lastly, signals were *Z*-standardized by shifting to zero mean and scaling to unit variance.

We subdivided each participant’s brain into 214 regions using the Schaefer atlas^190^ to represent 200 cortical regions assigned to 17 subnetworks and the Harvard Oxford atlas^191^ to represent 14 subcortical regions. For each of the 214 regions, an average BOLD time-series was computed across voxels. For the main network control theory analysis, each time-series was further separated by the thought and n-back block based on the timing information (i.e., onset and duration of each task block during fMRI scanning) extracted from raw E-Prime experimental logs.

The ACC includes the perigenual and what has also been named the midcingulate regions^192,193^. We used the bilateral parcels “LH_ContA_Cingm_1” and “RH_ContA_Cingm_1” in the Schaefer atlas. Approximately 43% of the voxels in these two parcels overlap with a prior dorsal ACC region of interest^194,195^.

### Statistical analysis

Between-group differences (clinical vs. non-clinical) were calculated using two-sample *t*-tests or Mann-Whitney U-tests. Within-group comparisons of neural measures during neutral thought and neural measures during rumination or worry were conducted using paired *t*-tests and Wilcoxon match paired tests.

Symptom severity was predicted using GLMs. In each model, the dependent variable was defined as one of the dimensional phenotypes of depression, anxiety, rumination, or worry. The independent variable was the neural metric of interest, including average transition probability, fractional occupancy, or control energy, with age and sex serving as covariates. We adjusted for multiple comparisons and report findings under FDR and Bonferroni correction, respectively.

### Dynamic brain states

We used a discrete model as a simplification of brain dynamics to characterize BOLD fMRI activity patterns. Repeatedly visited locations in regional activation space are neural representations of cognitive states, which are termed “brain states”. We used *k*-means clustering to characterize the progression of brain states over time.

First, we concatenated all functional volumes into one large data matrix. Then, to determine the brain states present in these data, we performed 20 repetitions of *k*-means clustering for *k* = 2 to *k* = 11 using Pearson correlation as the algorithm’s measure of distance. Because we aimed to study the temporal progression between coactivation patterns, we created a *k*-by-*k* transition probability matrix between clusters of coactivation patterns. Based on prior work applying this approach^63,133^, we selected *k* based on the lowest error solution across repetitions, the variance explained, and the change in variance explained for a unit increase in *k*. We quantified the quality of the partition by calculating the adjusted mutual information between iterations.

We observed that the variance explained by the clustering algorithm began to taper off after *k* = 6 and this is also where the average silhouette width was greatest, a metric that quantifies the quality of the clustering based on the separation of the discrete states. Additional brain states were not all represented in every participant, which would present a challenge to our statistical tests for cross-participant comparisons of state dynamics. To further validate the choice of *k* = 6, we evaluated the split-half reliability of the partition at this resolution which was *r* = 0.47. The reliability when splitting across the thought block portion of the task and the 1-back working memory task was *r* = 0.46.

After using *k*-means clustering to define discrete brain states, we generated names for each state using the maximum cosine similarity to binary vectors reflecting activation of communities in an *a priori*-defined 17-network partition^190^. Using the brain states, we calculated the transition probability and fractional occupancy of states. The transition probability between state *i* and state *j* is the probability that *j* is the next new state occupied after state *i*. This is the probability of a particular state transition occurring given a change in states. Fractional occupancy is the percentage of volumes in each scan that were classified as a particular state.

### Network control theory

The network control theory framework has been used to determine how underlying white matter architecture constrains transitions between different brain states inferred from neuroimaging data^65,71,130,196^. The control inputs required to execute these transitions between brain states can be considered a way of operationalizing top-down control signals involved in regulating perseverative thought. Importantly these control signals were simulated from known regions associated with top-down regulation^73,190,197,198^ and the magnitude of the signals were calculated from the combination of bi-directional activity across all connected brain regions over time.

More concretely, the dynamics were defined by the following continuous-time system equation based on procedures described in more detail in prior work^68^:

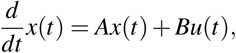

where 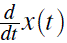 is the states of downstream nodes following the change to the states of the upstream nodes *x*(*t*) with control signals *u*(*t*), **A** is the 214-by-214 region connectome, and **B** is a 214-by-214 matrix where the diagonal entries contain the scalar values of 1 for the regions in the frontoparietal and attention networks sending control signals. Because we lacked diffusion imaging data, and to focus our analysis on spatiotemporal brain patterns, we used an average structural network template for all network control theory simulations obtained from prior work^73^ as **A**. We defined **B** to allow control inputs into the 145 brain regions of the dorsal attention, ventral attention, and frontoparietal networks, following prior cognitive neuroscience literature implicating these networks in exerting executive control resources^73,190,197,198^.

To determine the optimal control energy to transition from an initial state *x*_0_ to a target state *x _f_*, a cost function was optimized to minimize both control energy and the distance to target states for selected nodes over a finite time horizon. The unique control input *u*^∗^(*t*) needed to transition the system from an initial state *x*(0) = *x*_0_ to a final target state *x*(*T*) = *x _f_* over the time horizon *T* is determined through the cost function that solves the problem:

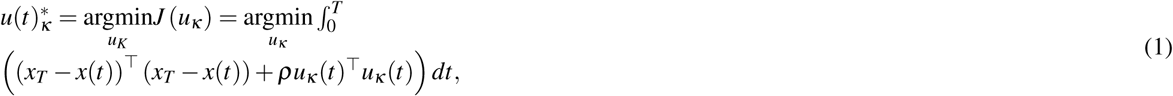

where the parameter *ρ* determines the relative weighting between the costs associated with the length of the state trajectory and the input energy. Following prior work, we set *ρ* to 1 such that no specific assumptions are made about the relative importance of constraints on energy and distance values^199^. The cost function 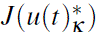 is defined to find the unique optimal control input 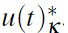. As in prior work^65,73^, this cost function was parameterized by *ρ* = 100 for a time horizon *T* = 3 to tune the mixture of these two costs while finding the input *u*(*t*) that achieves the state transition. We then use this optimal control input to calculate the control inputs required by a single brain region by integrating each control input over 1000 time steps in the simulation.

In our model, we used successive activity patterns (TRs) during the task time series at *x_t_* and *x_t_*_+1_ as the initial state *x*_0_ and the target state *x _f_*. Calculating each transition across successive TRs in each individual’s time series provides us with values of the control energy that were averaged within each thought and 1-back period. For the analysis relating control energy to the dynamic brain state measures of transition probability and fractional occupancy, we defined the states *x*_0_ and *x _f_* by the centroid of each of the 6 brain states detected by *k*-means clustering. For example, this approach allows us to correlate a 6-by-6 transition matrix with a 6-by-6 control energy matrix. Both regional and individual control energy values were used, where an individual’s energy value was defined by the average of their regional values.

## Acknowledgments

DZ acknowledges support from the George E. Hewitt Foundation for Medical Research. This research was supported by a University Research Foundation grant and a Center for Functional Neuroimaging Pilot Project Fund award to AMR, and by a MindCORE Collaborative Research grant to AMR and DSB.

## Supplementary Materials

**Figure S1.**
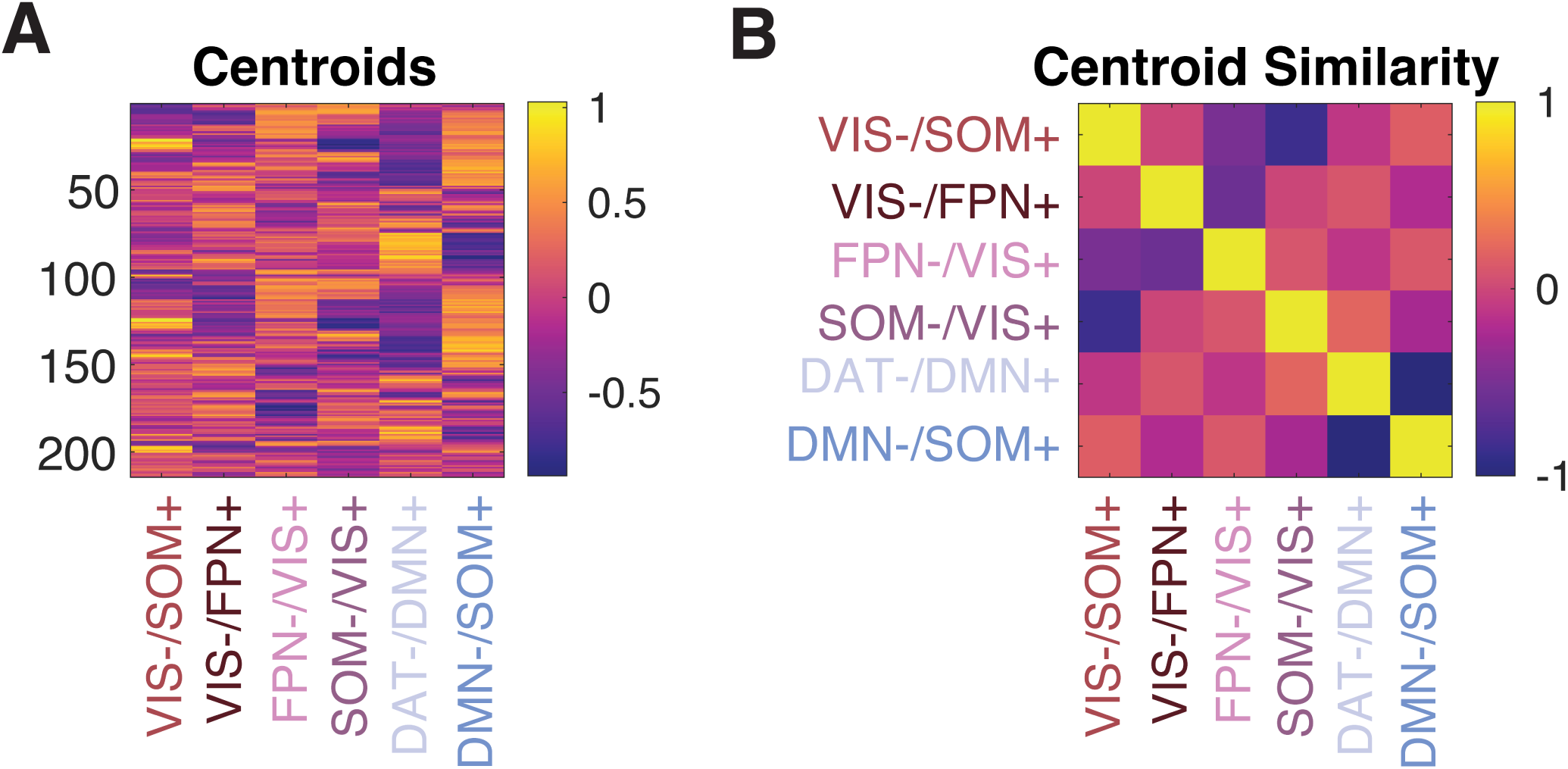
Brain states identified by *k*-means clustering. **(A)** A matrix of centroids depicting the fMRI activity across 342 repeated measurements (TRs) during the task clustered into six discrete brain states (columns) and their loading onto each brain region (rows). **(B)** A matrix of centroid similarity. Several states are anti-correlated, such as states with VIS– and VIS+ or DMN– and DMN+. Other states are relatively orthogonal to each other (centroid similarity ≈ 0).

**Figure S2.**
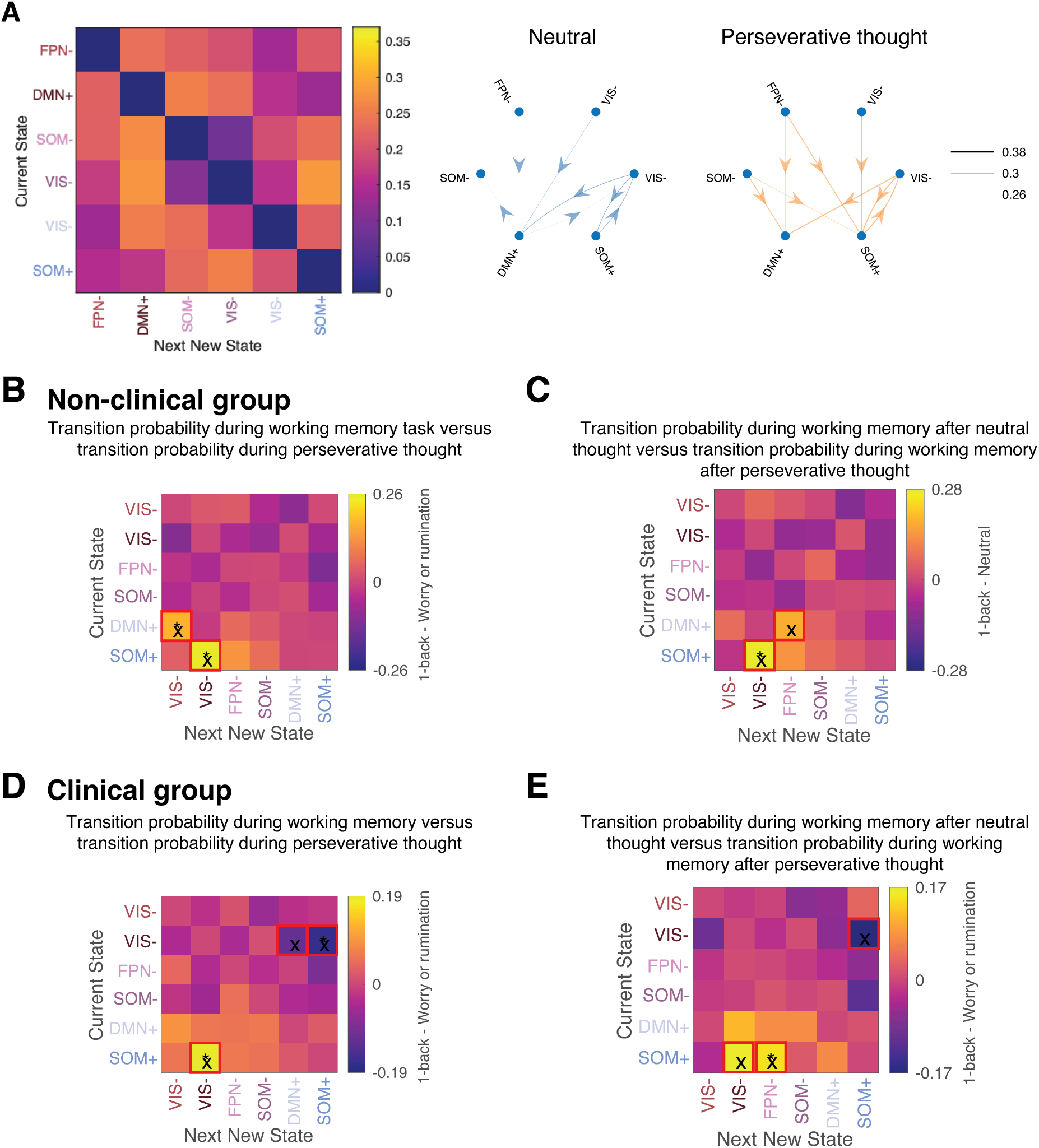
Transition probabilities differ during a working memory task compared to during perseverative thought. **(A)** *Left*: Probability of transitioning between the 6 brain states across the experiment. *Right*: Transitions visualized as a network. **(B)** Transition probabilities during perseverative thought and during 1-back performance within the non-clinical group. There was a greater probability of transitioning out of the DMN+ state into the VIS-/FPN+ state during 1-back performance than during worry or rumination. There was also a greater probability of transitioning from the SOM+ state into the VIS-/FPN+ state. **(C)** When the non-clinical group performed the 1-back task after worry or rumination, there was a higher probability of transitioning from DMN+ into FPN-as well as from SOM+/DMN-into VIS-/FPN+ than when the group performed the 1-back task after neutral thought. **(D)** Transition probabilities during perseverative thought and during 1-back performance within the clinical group. There was a reduced probability of transitioning into the DMN+ state from the VIS-/FPN+ state during the 1-back task than during worry or rumination. There was also a lower probability of transitioning out of the VIS-/FPN+ state into the SOM+/DMN-state as well as a greater probability of transitioning out of the SOM+/DMN-state into the VIS-/FPN+ state during the 1-back task than during perseverative thought. **(E)** When the clinical group performed the 1-back task after worry or rumination, there was a higher probability of transitioning from SOM+/DMN-into VIS-/FPN+ as well as from SOM+/DMN-into FPN-than when the group performed the 1-back task after neutral thought. X: FDR-corrected *p* < 0.05; *:

**Figure S3.**
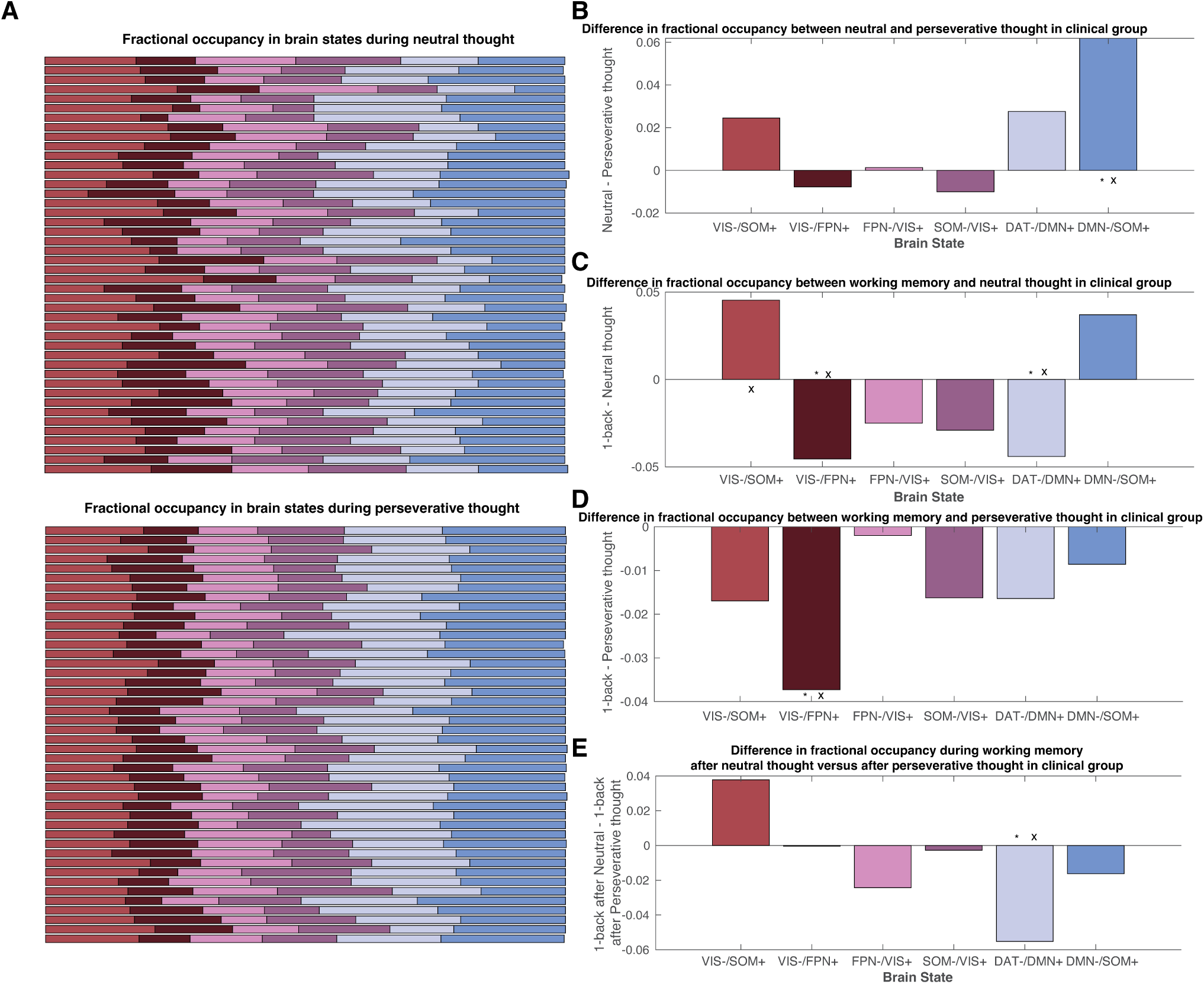
Fractional occupancies differ during neutral thought, perseverative thought, and working memory within the clinical group. **(A)** Fractional occupancy in the six brain states for each participant (rows) during neutral thought (*top*) and during perseverative thought (*bottom*). **(B)** The clinical group had greater fractional occupancy of the SOM+/DMN-brain state during neutral thought than during perseverative thought. **(C)** The clinical group had greater fractional occupancy in DAT-/DMN+ during neutral thought than during performance of the 1-back task. This group also had greater fractional occupancy of the VIS-/FPN+ brain state during the 1-back task than during neutral thought. **(D)** The clinical group had greater fractional occupancy in the VIS-/FPN+ brain state during perseverative thought than during the 1-back task. **(E)** The clinical group dwelled in the DAT-/DMN+ state more during the 1-back task when the 1-back task followed perseverative thought than when it followed neutral thought. In panels B-E, X represents FDR-corrected *p* < 0.05, whereas * represents Bonferroni-corrected *p* < 0.05.

**Figure S4.**
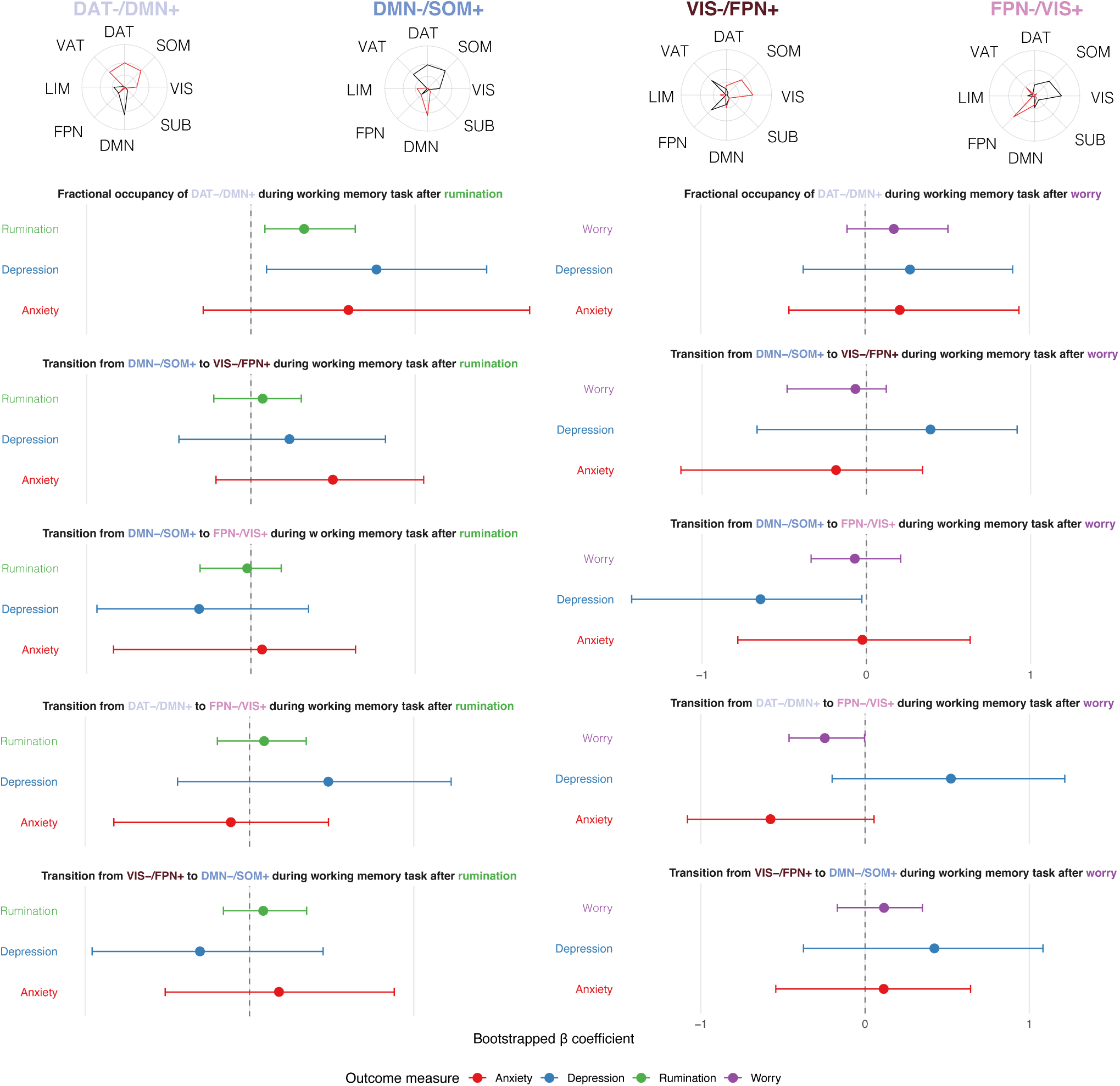
Relationship between dynamic neural metrics and behavioral phenotypes. *Top*: Radar plots of neural states of interest involving the frontoparietal and default mode networks. *Bottom*: Linear regression models testing the relationships between a dynamic neural measure—during the rumination period (left column) or the worry period (right column)—and trait rumination, trait worry, major depression severity, and generalized anxiety severity while controlling for age and sex. Brackets depict 95% confidence intervals bootstrapped over 1000 iterations. Greater fractional occupancy of DAT-/DMN+ during the 1-back task when it followed rumination was associated with greater trait rumination severity. No effects were observed for worry.

**Figure S5.**
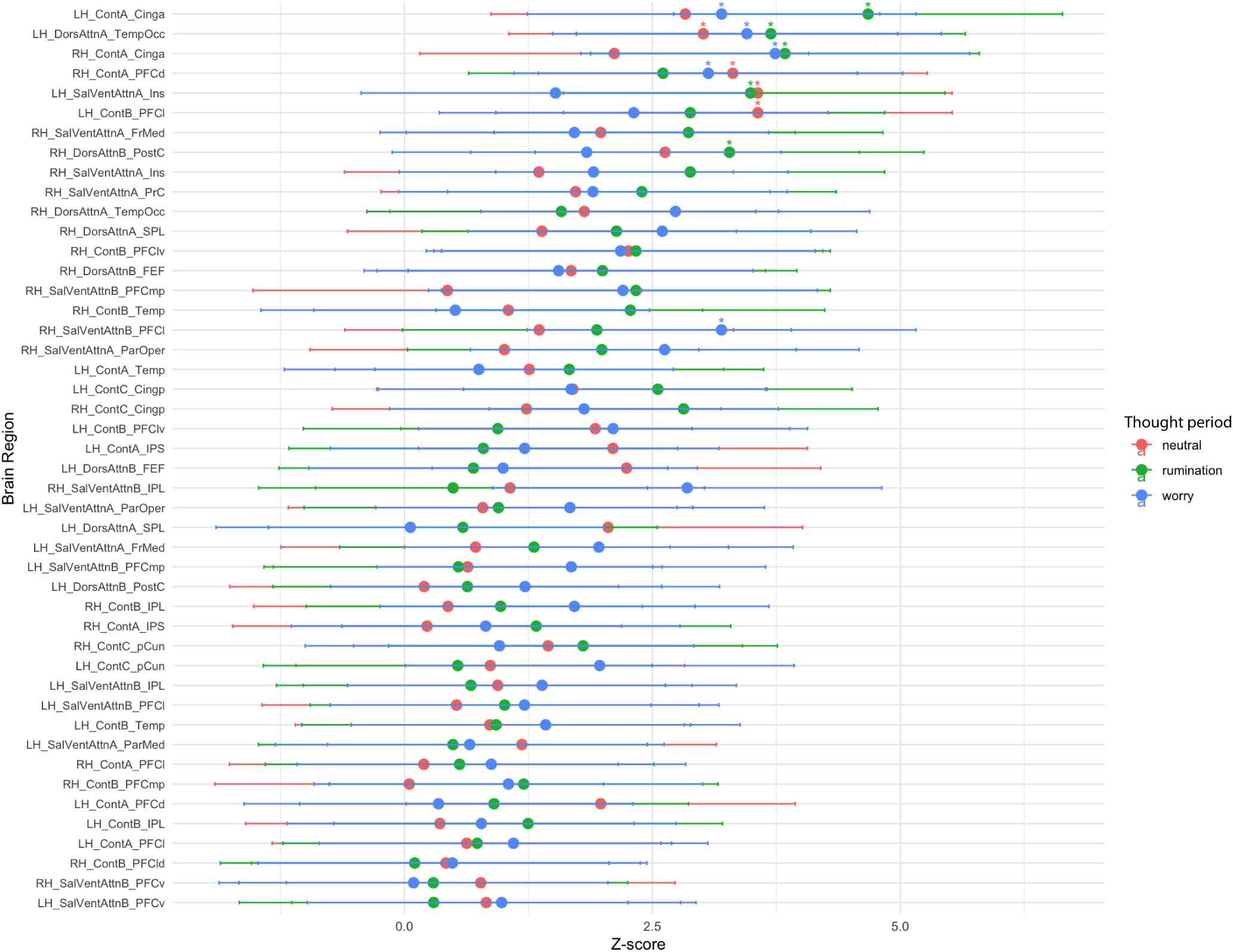
Regional control energy differences between clinical and non-clinical groups during different thought periods. *Z*-scores that are more positive indicate regions where the non-clinical group used more energy than the clinical group during the neutral (red), rumination (green), and worry (blue) thought periods. Asterisks indicate significant differences in control energy between groups (*p <* 0.05) after adjusting for multiple comparisons using FDR correction.

**Figure S6.**
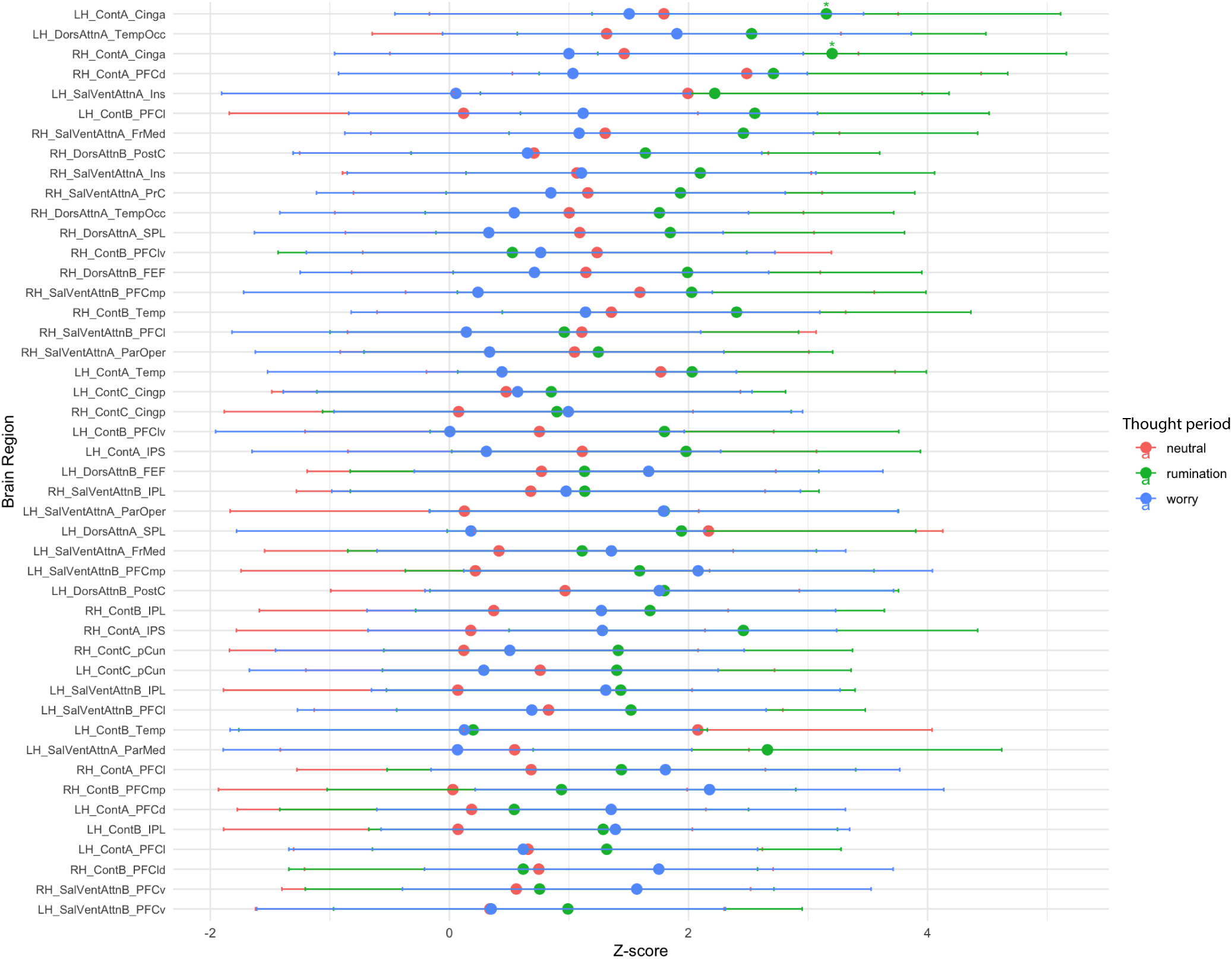
Regional control energy differences between clinical and control groups during the working memory task after different thought periods. *Z*-scores that are more positive indicate regions where the non-clinical group used more energy than the clinical group. Control energy differences between groups during the 1-back task when the task followed the neutral (red), rumination (green), and worry (blue) thought periods. Asterisks indicate significant differences in control energy between groups (*p <* 0.05) after adjusting for multiple comparisons using FDR correction.

## Citation Diversity Statement

Recent work in several fields of science has identified a bias in citation practices such that papers from women and other minority scholars are under-cited relative to the number of such papers in the field^200–208^. Here we sought to proactively consider choosing references that reflect the diversity of the field in thought, form of contribution, gender, race, ethnicity, and other factors. First, we obtained the predicted gender of the first and last author of each reference by using databases that store the probability of a first name being carried by a woman^204,209^. By this measure (and excluding self-citations to the first and last authors of our current paper), our references contain 17.03% woman(first)/woman(last), 15.38% man/woman, 17.29% woman/man, and 50.29% man/man. This method is limited in that a) names, pronouns, and social media profiles used to construct the databases may not, in every case, be indicative of gender identity and b) it cannot account for intersex, non-binary, or transgender people. Second, we obtained predicted racial/ethnic category of the first and last author of each reference by databases that store the probability of a first and last name being carried by an author of color^210,211^. By this measure (and excluding self-citations), our references contain 7.8% author of color (first)/author of color(last), 15.33% white author/author of color, 18.63% author of color/white author, and 58.24% white author/white author. This method is limited in that a) names and Florida Voter Data to make the predictions may not be indicative of racial/ethnic identity, and b) it cannot account for Indigenous and mixed-race authors, or those who may face differential biases due to the ambiguous racialization or ethnicization of their names. We look forward to future work that could help us to better understand how to support equitable practices in science.

